# SIMBSIG: Similarity search and clustering for biobank-scale data

**DOI:** 10.1101/2022.09.22.509063

**Authors:** Michael F. Adamer, Eljas Roellin, Lucie Bourguignon, Karsten Borgwardt

## Abstract

**Summary:** In many modern bioinformatics applications, such as statistical genetics, or single-cell analysis, one frequently encounters datasets which are orders of magnitude too large for conventional in-memory analysis. To tackle this challenge, we introduce SIMBSIG, a highly scalable Python package which provides a scikit-learn-like interface for out-of-core, GPU-enabled similarity searches, principal component analysis, and clustering. Due to the PyTorch backend it is highly modular and particularly tailored to many data types with a particular focus on biobank data analysis.

**Availability:** SIMBSIG is freely available from PyPI and its source code and documentation can be found on GitHub (https://github.com/BorgwardtLab/simbsig) under a BSD-3 license.

## 1 Introduction

With the ever increasing amount of data produced, e.g. by genomics and next generation sequencing, there is also demand for powerful computational tools to analyse such state-of-the-art datasets. In the UK Biobank alone, genotype data (800, 000 single nucleotide polymorphisms, SNPs) of half a million people is available (Bycroft *et al*., 2018). This genotype dataset vastly exceeds the size of random access memory (RAM) or graphics processing units (GPU) memory (VRAM) of conventional computers. Therefore, algorithms and data formats, such as hdf5 files, which allow for efficient out-of-core computations are needed to perform any analysis or data processing. In order to tackle this issue, we introduce “SIMBSIG = <u>SIM</u>milarity <u>B</u>atched <u>S</u>earch Integrated <u>G</u>PU”, which can efficiently perform nearest neighbour (NN) searches, principal component analysis (PCA), and k-means clustering on central processing units (CPUs) and GPUs, both in-core and out-of-core. SIMBSIG bridges the gap between the functionality and ease-of-use of the popular scikit-learn package and the speed of current computer hardware.

While there are many packages with similar functionality to SIMBSIG, most noteworthy, the Faiss software (Johnson *et al*., 2019), the lack of a scikit-learn-like interface and the relatively rigid set of distance metrics make it a less tailored to bioinformatics applications. Nevertheless, a future version of SIMBSIG might incorporate Faiss as an alternative backend to PyTorch.

## 2 Features

SIMBSIG is a Python package which utilises PyTorch to ensure efficient computations and provides a high-level interface mimicking scikitlearn. At its heart, SIMBSIG comprises three modules, `neighbors, clustering`, and decomposition. All methods accept numpy arrays or hdf5 file handles as inputs. Due to the modular structure of SIMBSIG, it will be straightforward to extend it to other data types.

We chose numpy arrays as one of our input types for the user to have a seamless transition from scikit-learn while being able to use GPU acceleration whenever needed. The `hdf5` format is a hierarchical data format and provides many advantages over other tabular-data formats, which makes `hdf5` file objects ideal inputs to SIMBSIG. GPU usage with SIMBSIG is optional, enabling users to apply the full flexibility offered by SIMBSIG, even without access to GPUs.

**Fig. 1.**
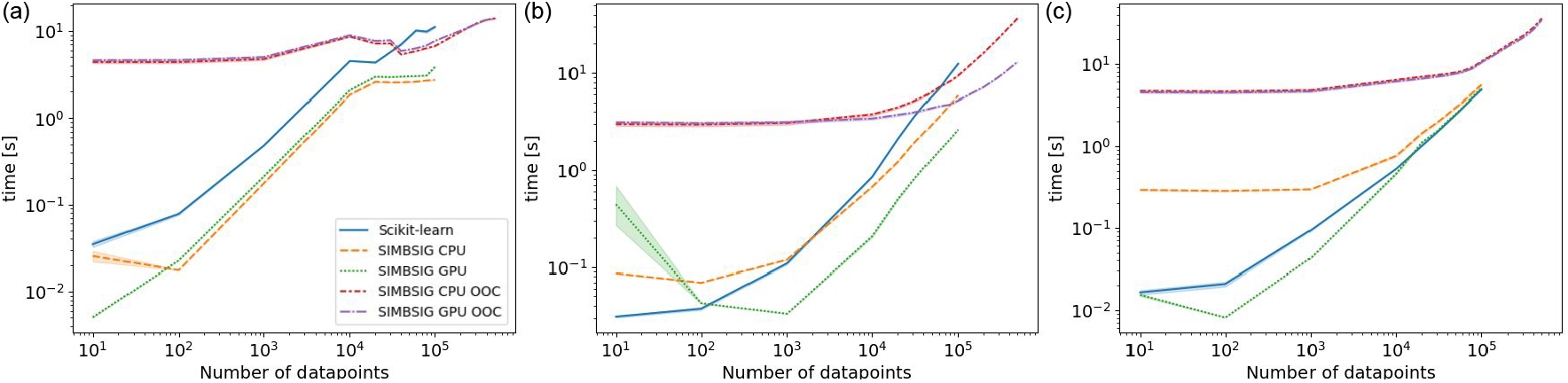
The runtime performance of SIMBSIG compared to the scikit-learn baseline averaged over 10 runs. We tested KMeans (a), a KNN search (b), and PCA (c). Both, in-core and out-of-core (ooc) scenarios were computed, whereas the maximum dataset size for in-core was 10^5^ datapoints. The batch size was 10^4^ in order to maximally use the available VRAM. All runs were performed on an Intel i7 10700K CPU and 32GB RAM, the GPU was an Nvidia GeForce RTX3080 with 10GB VRAM. Stopping criteria for KMeans were set to default values.

The `neighbors` module consists of our core routine for nearestneighbours searches in a batched, brute force manner. This guarantees exact results with a precisely controllable memory-runtime trade-off. We implemented a batched K-Nearest Neighbors (KNN) search, where the number K of neighbors is fixed a priori, and a radius neighbour search, where all neighbours within a user-defined radius are returned. Based on this core routine, we implemented classifiers and regressors.

We use a brute-force approach only due to the infeasibility of other exact methods in this scenario, while retaining most other functionality of scikit-learn such as the choice of a range of metrics including all *l*_*p*_ distances. Since, especially in genetic applications, the data is often high dimensional and subject to the curse of dimensionality, we also implemented the fractional *l*_*p*_ distance (Aggarwal *et al*., 2001). We further allow for precomputed distance matrices or user-defined functions (“callables”). The “callable” functionality is also present in scikit-learn and provides an easy way for more sophisticated (dis-)similarity functions such as kernel functions. Kernels exist, for example, for SNP data (Wessel and Schork, 2006) and may help obtaining more accurate results.

The second core routine is batched k-means clustering from the `clustering module`. The implementation is based on the fast k-means algorithm by Sculley (2010). Similarly to the NN search, we accept numpy arrays or hdf5 file handles as input and we can cluster with respect to any *l*_*p*_ norm, the cosine distance or a “callable” distance function. We additionally implemented a heuristic stopping criterion based on the maximum relative change of any cluster centre. In practice, this stopping criterion works well for the *l*_*p*_ distances, however, it may be lacking an analogue for more involved distance functions. In the current implementation, for any metrics that are not *l*_*p*_, the relative change becomes an absolute change.

The third pillar of SIMBSIG is the `decomposition` module. This module provides an out-of-core, GPU-accelerated implementation of Halko’s approximate PCA algorithm (Halko *et al*., 2011)

## 3 Experiments

To test accuracy, we designed an extensive set of unit tests, where we compare all possible input choices to the output of scikit-learn which we assume to be the ground truth.

The speed of SIMBSIG was benchmarked on a large, artificial SNP dataset, where SNPs are encoded according to dominance assumption. We sampled “participants” represented by a 10, 000 dimensional vector with independent entries in *{*0, 1, 2*}*, representing 10, 000 SNPs with probabilities *{*0.6, 0.2, 0.2*}*. In total, we sampled 500, 000 such participants and stored the dataset in the hdf5 format. We tested in-memory CPU performance against scikit-learn and benchmarked outof-core scenarios. For in-core computations, GPU accelerated SIMBSIG similarity searches are ∼ 1 order of magnitude faster compared to scikit-learn. Out-of-core similarity searches run both with and without GPU acceleration, where SIMBSIG’s GPU acceleration feature offers a significant reduction in execution time for these heavy computations. A reference GPU implementation given by cuML (Raschka *et al*., 2020) could not use the `h5py` package needed to load hdf5 files.

## Funding

This work was supported by the Swiss National Science Foundation [# PZ00P3_186101]; and the Alfried Krupp Prize for Young University Teachers of the Alfried Krupp von Bohlen und Halbach-Stiftung [K.B.]

## Notes

### Competing Interest Statement

The authors have declared no competing interest.

## References

Aggarwal, C. C. et al. (2001). On the surprising behavior of distance metrics in high dimensional space. In International conference on database theory, pages 420–434. Springer.

Bycroft, C. et al. (2018). The UK biobank resource with deep phenotyping and genomic data. Nature, 562(7726), 203–209.

Halko, N. et al. (2011). An algorithm for the principal component analysis of large data sets. SIAM Journal on Scientific computing, 33(5), 2580– 2594.

Johnson, J. et al. (2019). Billion-scale similarity search with GPUs. IEEE Transactions on Big Data, 7(3), 535–547.

Raschka, S. et al. (2020). Machine learning in python: Main developments and technology trends in data science, machine learning, and artificial intelligence. arXiv preprint arXiv:2002.04803.

Sculley, D. (2010). Web-scale k-means clustering. In Proceedings of the 19th international conference on World wide web, pages 1177–1178.

Wessel, J. and Schork, N. J. (2006). Generalized genomic distance– based regression methodology for multilocus association analysis. The American Journal of Human Genetics, 79(5), 792–806.

